# Uncovering Predictive Gene and Cellular Signatures for Checkpoint Immunotherapy Response through Machine Learning Analysis of Immune Single-Cell RNA-seq Data

**DOI:** 10.1101/2024.11.16.623986

**Authors:** Asaf Pinhasi, Keren Yizhak

## Abstract

**Background:** Immune checkpoint inhibitors have revolutionized cancer therapy by harnessing the body’s immune system to eliminate tumor cells. However, only a subset of patients responds to treatment. Understanding why only some patients respond remains a critical challenge in cancer research due to the complexity and variability of cellular composition within the tumor microenvironment. Our study presents PRECISE (*Predicting therapy Response through Extraction of Cells and genes from Immune Single-cell Expression data*), a pipeline that leverages single-cell RNA sequencing data and machine learning techniques to predict ICI responses, while maintaining the richness of single-cell information and ensuring interpretability of the results.

**Results:** The PRECISE pipeline implements gene and cell-filtering approaches for optimizing treatment prediction. Utilizing the XGBoost algorithm for predicting patient response to ICI on a dataset of melanoma-infiltrated immune cell achieved an initial Area Under the Curve (AUC) of 0.84. This signal is further improved to 0.89 after Boruta feature selection, revealing an 11-gene predictive signature. Investigation of these genes through SHAP values identified various gene-pair interactions with non-linear conditional effects on predictions. Furthermore, a novel reinforcement learning framework implemented in PRECISE reveals non-predictive single cells that detracts the model’s performance. Altogether, the identified gene- and cell-based signatures demonstrates high prediction power across independent datasets, including lung, breast, brain, and skin cancers.

**Conclusions:** Our approach demonstrates the potential of supervised machine learning and reinforcement learning to enhance the understanding of cancer immunity and improve the prediction of treatment responses using single-cell data.

Understanding patients’ response to immune checkpoint inhibitors (ICIs) is a critical challenge in cancer research, given the complexity and variability of immune interactions within the tumor microenvironment. Our study leverages single-cell RNA sequencing data and aims to utilize machine learning techniques to predict immunotherapy responses, while maintaining the richness of single-cell information and ensuring interpretability of the results. Using a dataset of melanoma immune cells, we conducted thorough preprocessing and applied the XGBoost algorithm in a leave-one-out cross-validation fashion to predict sample response. By labeling cells according to their sample’s response and aggregating predictions, we achieved an initial AUC score of 0.84. This score was improved to 0.89 with the application of Boruta feature selection, which identified key predictive genes, leading to an 11-gene predictive signature. Further analysis pinpointed T cell clusters as significant contributors to immune response. Utilizing SHAP values provided deeper insights into gene behaviors, interactions, and their effects on the model’s predictions. Additionally, a novel reinforcement learning model was developed for single-cell level prediction and characterization, allowing us to identify and analyze the most predictive cells for response and non-response. Our approach demonstrates the potential of sophisticated computational methods to enhance our understanding of cancer immunity and improve the prediction of treatment responses.

## Introduction

Cancer immunotherapy, particularly immune checkpoint inhibitors (ICI), has revolutionized cancer treatment, offering durable responses in a subset of patients. ICI, such as anti-PD1 antibodies, work by reinvigorating exhausted T cells within the tumor microenvironment (TME), allowing the immune system to act effectively against tumor cells (1). Despite their success, the majority of patients fail to respond to these therapies (2), underscoring the need for reliable predictive biomarkers to identify patients who will likely benefit from treatment. Many efforts aim to decipher the behavior of the immune system against the tumor as a result of treatment (3). In particular, characterizing the immune system in relation to different ICI responses and finding robust biomarkers to differentiate between responders and non-responders has become a major goal in the field of cancer immunity, often focusing on different cells in the TME (4–6). Nevertheless, the complexity of immune interactions, and the context dependent behavior of the immune system to treatment are main obstacles in the endeavor for enhancing the efficacy of immunotherapy treatments.

Single-cell RNA sequencing (scRNA-seq) provides unparalleled insights into the cellular heterogeneity and dynamic states within the TME, capturing the complexity of immune responses at a granular level. However, despite the huge advances in the field of machine learning (ML), integrating this high-dimensional data with advanced models and techniques to predict therapeutic outcomes remains a challenge. Most existing approaches either oversimplify the data, losing valuable single-cell resolution (7,8), or fail to provide interpretable results, limiting their clinical applicability (9–11). Bridging this gap requires the development of innovative computational methods that retain data richness while offering clear insights into the biology underpinning the decision making of the models.

While machine learning in general has been widely adopted in the field of single-cell analysis (12), relatively few studies use supervised learning to directly predict response to immunotherapy. For example, Dong et al. used cell-type proportion extracted from scRNA-seq data for training ML models to predict response to ICI (8). In a different study (7), Kang et al. used pseudo-bulk expression calculated from scRNA-seq. Prediction models with similar aims to ours have been developed, such as CloudPred (13), which uses differentiable machine learning to predict lupus, and ScRAT (14), which uses neural networks to predict COVID-19 phenotypes. However, these models have not been applied in the context of immunotherapy response.

Applying ML to scRNA-seq data raises a few challenges and open questions: for instance, should all single cells from a responding patient be labeled as ‘responders’, or should we use pre-existing knowledge on cell function to label cells as either ‘‘favorable’ or ‘unfavorable’? How can we infer the patient class from the predicted classes of the associated single cells? Should we use all cells for prediction or perhaps a subset that is more predictive than others?

In this study we addressed these various questions by investigating scRNA-seq data of *CD45*^+^ cells from a cohort of ICI-treated melanoma patients (15). We developed **PRECISE** (*Predicting therapy Response through Extraction of Cells and genes from Immune Single-cell Expression data*), a machine learning framework designed to predict immunotherapy response by and extract gene- and cell-level insights from scRNA-seq data. Using XGBoost (eXtreme Gradient Boosting) as the ML model (16), our basic methodology involved labeling cells based on their sample’s response, training models in a leave-one-out manner, and then aggregating predictions to create a sample-level score. For feature selection (FS) we used Boruta (17) which provided high performance in predicting patient response (AUC = 0.89). Following, we dissected the contribution of selected genes using feature importance scores and the more advanced Shapley Additive exPlanations (SHAP) values (18), identifying complex, non-linear contribution of single genes and gene pairs to treatment response. Considering the cells-axis, we identified specific cell-types that are more informative than others in predicting response. Moreover, we devised a novel reinforcement learning model designed to identify single cells that are either predictive or non-predictive of patient response (19). Applying these gene- and cell-based signatures to different datasets, we managed to achieve high prediction power across different cancer types. Overall, our findings highlight the potential of advanced computational approaches in enhancing our understanding of cancer immunity and informing treatment.

## Results

### Base Model Feature Selection and Per Cell Type Prediction

We first utilized our previously published (15) single-cell RNA sequencing (scRNA-seq) dataset that consists of 16,291 CD45^+^ cells from 48 tumor biopsies taken from 32 stage IV metastatic melanoma patients treated with ICI. 11 patients had longitudinal biopsies, and 20 patients had one or two biopsies taken either at baseline or during treatment. Tumor samples were classified according to radiologic evaluations as either progression/non-responder (NR; n = 31, including stable disease (SD) and progressive disease (PD) samples) or regression/responder (R; n = 17, including complete response (CR) and partial response (PR) samples). After the removal of non-informative genes (Methods), we constructed an XGBoost-based machine-learning model and investigated its power in predicting the patients’ response to ICI (Figure 1A).

**Figure 1:**
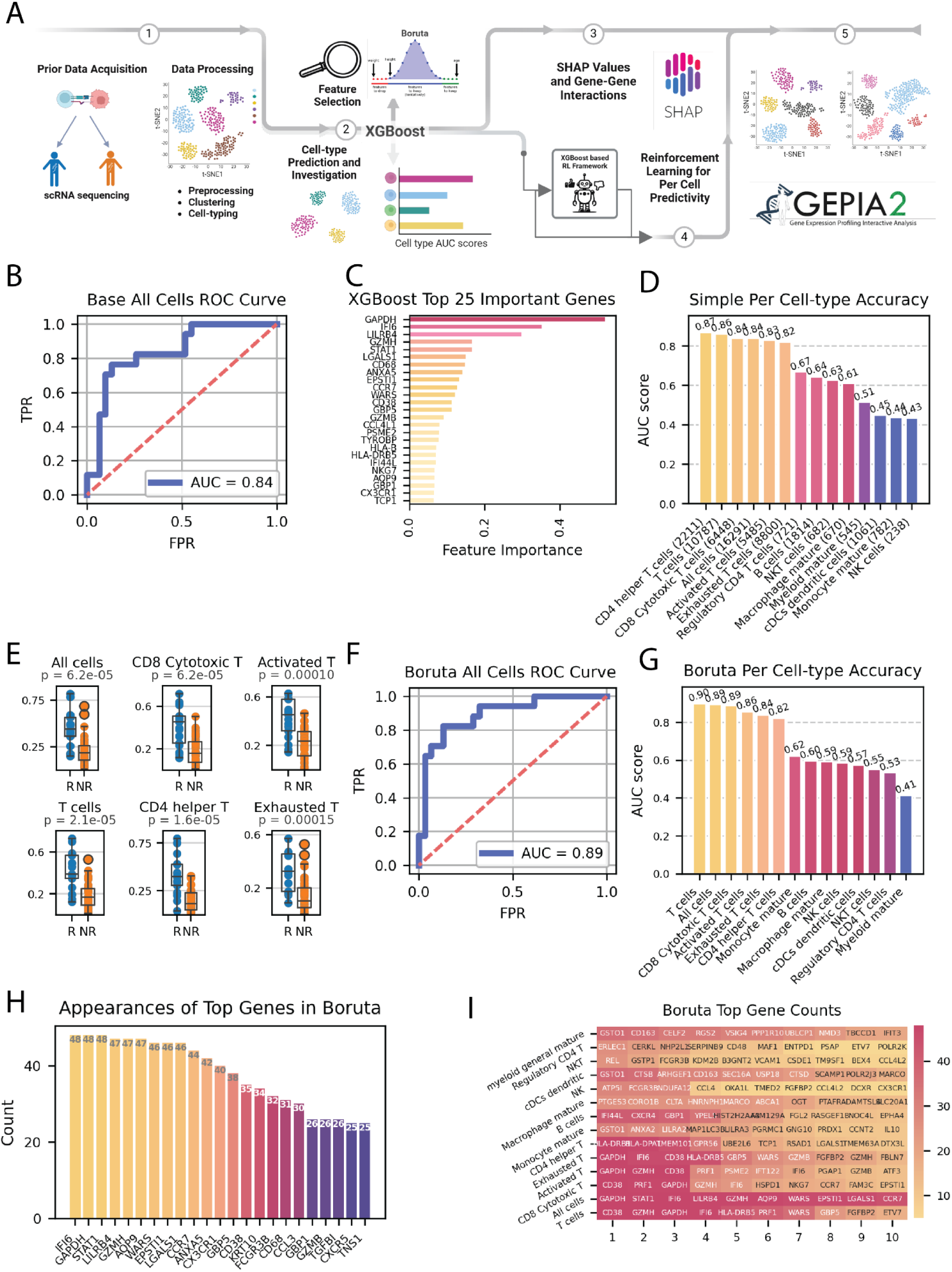
Prediction of ICI Response Using XGBoost and Boruta Feature Selection per Cell-Type. **A** Schematic workflow of the study. **B** ROC curve for the base model predicting response to ICI using all cells and genes in the cohort. **C** XGBoost feature importance bar-plot showing the top 30 most important genes. **D** AUC scores indicating the prediction accuracy of the base model across different immune cell types. **E** Box plots comparing the scores produced by base XGBoost between responders (R) and non-responders (NR) across top most accurate cell subtypes, with significant differences indicated by the Mann–Whitney U test. **F** AUC scores indicating the prediction accuracy of the Boruta selected model across different immune cell types. **G** ROC curve for the Boruta-selected model predicting response to ICI using all cells in the cohort. **H** Bar-plot of the number of occurrences of each gene in the Boruta selection across the LOO folds, showing the top most robust genes. **I** Heatmap displaying the top genes selected by Boruta for different immune cell types, showing number of occurrences of each gene for each cell type.

One of the main challenges in using supervised machine learning with single-cell data is the imbalance between the vast amounts of single-cells and the typically small number of samples from which these cells are taken. Naturally, the prediction target is often at the sample/patient level, raising the question of how best to assign labels to individual cells. To address this, we adopted a two-step approach: we trained the model and made predictions at the single-cell level, and then aggregated these predictions to devise a sample score. Each cell was labeled according to the response status of its sample of origin, such that cells from responding samples were classified as ‘responders’ and those from non-responding samples as ‘non-responders’. After predicting the labels for all cells in a sample, we calculated the proportion of cells predicted as ‘responders,’ and this proportion constituted the sample score. The logic for this scoring could be described as a tug-of-war between ‘good’ and ‘bad’ cells. The more ‘good’ cells a sample contains, the more likely it is to be pulled towards a favorable response. Considering the small number of samples, training and testing was conducted in a leave-one-out (LOO) cross-validation manner, with each fold training the model on the cells from all samples except one, which was excluded for prediction. This approach also mitigates the stochastic effects often encountered in standard cross-validation.

As our reference model, we included all 16,291 cells and all 10,082 genes that passed quality control (Methods). The ROC AUC score for this analysis was 0.84, showing reasonably high predictive accuracy (Figure 1B). To identify genes that are important for predicting patient response to ICI, and can therefore serve as potential biomarkers, we used the feature importance metric, which measures the contribution of a feature to the prediction of the model based on how often it was used to split the data in the ensemble of trees. To quantify the importance of each gene in the cohort we extracted the importance score from each of the 48 folds in the LOO simulation, and averaged them across all folds (Figure 1C). This analysis highlighted several genes as important for predicting patient response, including *GAPDH*, *IFI6*, *LILRB4*, *GZMH*, *STAT1*, *LGALS1* and *CD68* (Supplementary Table 2), all of which have already been found to relate to ICI and tumor immunity (20–25). For example, *CD68*, a marker for macrophages, was found to be associated with poorer clinical outcomes in multiple cancer types in a pan-cancer analysis of expression data from TCGA and GTEx (20). However, it is generally linked to M1 macrophages, which promote anti-tumor immunity (26). *STAT1* was found to promote immunotherapy response in melanoma (21), and was found to be positively correlated with *PD-L1* expression in ovarian cancer (22). *LILRB4*, has been found to strongly suppress tumor immunity, with studies in murine tumor models demonstrating that its blockade can alleviate this suppression, resulting in antitumor efficacy (23). Finally, *GAPDH*, receiving the highest feature importance score, has been shown to be involved in cancer progression and immune response regulation, particularly through its role in hypoxia-related pathways (24,25).

Examining the sample scores revealed that while most samples received scores consistent with their response status, some samples were mis-scored. For example, sample Pre_P3, a non-responder, received a high score, whereas sample Pre_P28, a responder, received a low score (Supplementary Table 1). These discrepancies could be due to model performance, but they might also indicate specific traits of these samples that warrant further investigation. Specifically, sample Pre_P3 had a mutation in *B2M* limiting antigen presentation, demonstrating how factors not shown in the transcription are causing the tumors to respond or resist to treatment. In addition, sample Pre_P28 was unique in its high abundance of cells from the myeloid lineage, emphasizing the importance of the specific cells used for prediction. Following, and after establishing the prediction power of the base model that considers all genes and all cells, we investigated the model’s performance when using specific subsets of these entities.

### Improving model performance by exploring cell and gene subsets

To improve model performance and study the contribution of different cell types and states, we first assigned individual cells to their respective cell type. We used the classification from the original paper which included 11 clusters determined by unsupervised clustering, and 13 cell types or states determined in a supervised manner using known gene markers (15). We then applied the XGBoost model to each of the different cellular groups separately. As expected, the per cell-type/cluster prediction resulted in different ROC AUC scores for each of the groups tested, testifying for their overall importance in predicting patient response. The results (Figure 1D, E) show that T cells subsets achieve substantially higher scores in comparison to the rest of the cell-types (0.87-0.82), in concordant with their known role in immune response in general, and specifically in melanoma (27). B cells (0.64), Macrophages (0.61) and other cell types were significantly less predictive. While the quantity of cells in each cellular groups is different and can potentially affect the accuracy of the model, this does not seem to be the dominant factor since there are T cell subsets that have a comparable number of cells to other cell-types, but produce much higher AUC scores (Figure 1D). Intriguingly, the model’s accuracy was consistently higher for the supervised cell types compared to the unsupervised clusters (Supplementary Figure 2), suggesting that the supervised clusters may capture more biologically meaningful signals and thus exhibit stronger predictive power.

We next moved to investigate if and to what extent feature selection would improve the base model predictions. To this end we explored 5 main methods for evaluating feature contribution - top highly variable (HVG), top differentially expressed, Lasso regression-based FS, simple top feature importance (Supplementary Table 3), and Boruta (Methods). Importantly, in all feature selection techniques applied, except for HVG method which is independent of response, we assured that selection was made independently from the predicted sample, thus avoiding any train-test leakage.

One of the main advantages of Boruta over all other feature selection techniques is the automatic determination of the number of features (17). While it is possible to tune Boruta hyperparameters to change the number of features, the method works well without any tuning, and offers robust feature number instead of an arbitrary choice. Therefore, although other feature selection techniques could offer slightly better accuracy for a specific choice of gene number or hyperparameters, Boruta was the best method for consistent and robust performance. The AUC scores produced by the other methods were mostly in the range 0.81-0.85, depending on the method and the hyperparameter, while Boruta produced a score of 0.89 (Figure 1F), manifesting its superiority in this selection.

Finally, to account for the observed variability in performance achieved for different cellular groups and gene subsets, we used Boruta feature selection to extract the top most contributing features on the base model and per cell-type, getting insight into the most important cell type-specific genes (Supplementary Table 4). To this end, we ran Boruta in the same manner as before, but this time focusing on each cell-type and cluster. As expected, the accuracy across most cell-types increased (Figure 1D,G). Exploring the top genes in each group (Figure 1H,I), we found genes with an established relation to cancer immunity and activation. For example, In NK cells, *FCGR3B*, known as *CD16b*, is involved in antibody-dependent cellular cytotoxicity (ADCC) (28,29). It was suggested that NK cells mediate ADCC-driven cytotoxicity specifically against *PD-L1*-positive cells, demonstrating the role of ADCC in enhancing the effectiveness of immunotherapies, particularly those targeting *PD-L1* (30). In B cells, *CXCR4* is expressed on the surface of mature B cells and interacts with *CXCL12* to recruit regulatory B cells to the tumor site. These cells can inhibit T cell activity, contributing to an immunosuppressive tumor environment. *HLA-DRB1* and *HLA-DP1*, both part of the MHC-II complex, were found in *CD4*^+^ helper T cells to be of the greatest importance. In addition, *CD38*^+^*HLA-DR*^+^ expressing *CD4*^+^ and *CD8*^+^ T cells were significantly expanded following ICI (immune checkpoint inhibitor) treatment in HNSCC (head and neck squamous cell carcinoma) (31). *GAPDH*, *CD38*, *STAT1* and *IFI6* all have well established roles in T cells activation and immunotherapy, and therefore they appear robust in most T cells subtypes (24,25,32–35). In addition to these general markers, more specific markers were extracted, such as *PRF1* in cytotoxic *CD8* T cell, a key component of the cytotoxic machinery of activated T cells, directly involved in killing tumor cells (36).

### SHAP Value Investigation Reveals Complex Gene Behaviors and Interactions

While XGBoost’s feature importance provides some insight into the most contributing genes, its interpretability is limited. For instance, feature importance is non-directional, and thus does not indicate whether a gene is more associated positively or negatively with response. To address this, and considering that gene behavior may extend beyond simple linear or dichotomous relationships, we employed SHAP (SHapley Additive exPlanations) values (18) (Methods). SHAP values, derived from cooperative game theory, measure both the direction and magnitude of a gene’s effect on the predicted response (Figure 2 A,B), allowing visualization of how gene expression influences it. Moreover, one of the strengths of SHAP values derives from its ability to capture context dependent contributions of each gene, such that the SHAP value for a gene can differ between two cells even if they share similar expression levels (Figure 2C).

**Figure 2:**
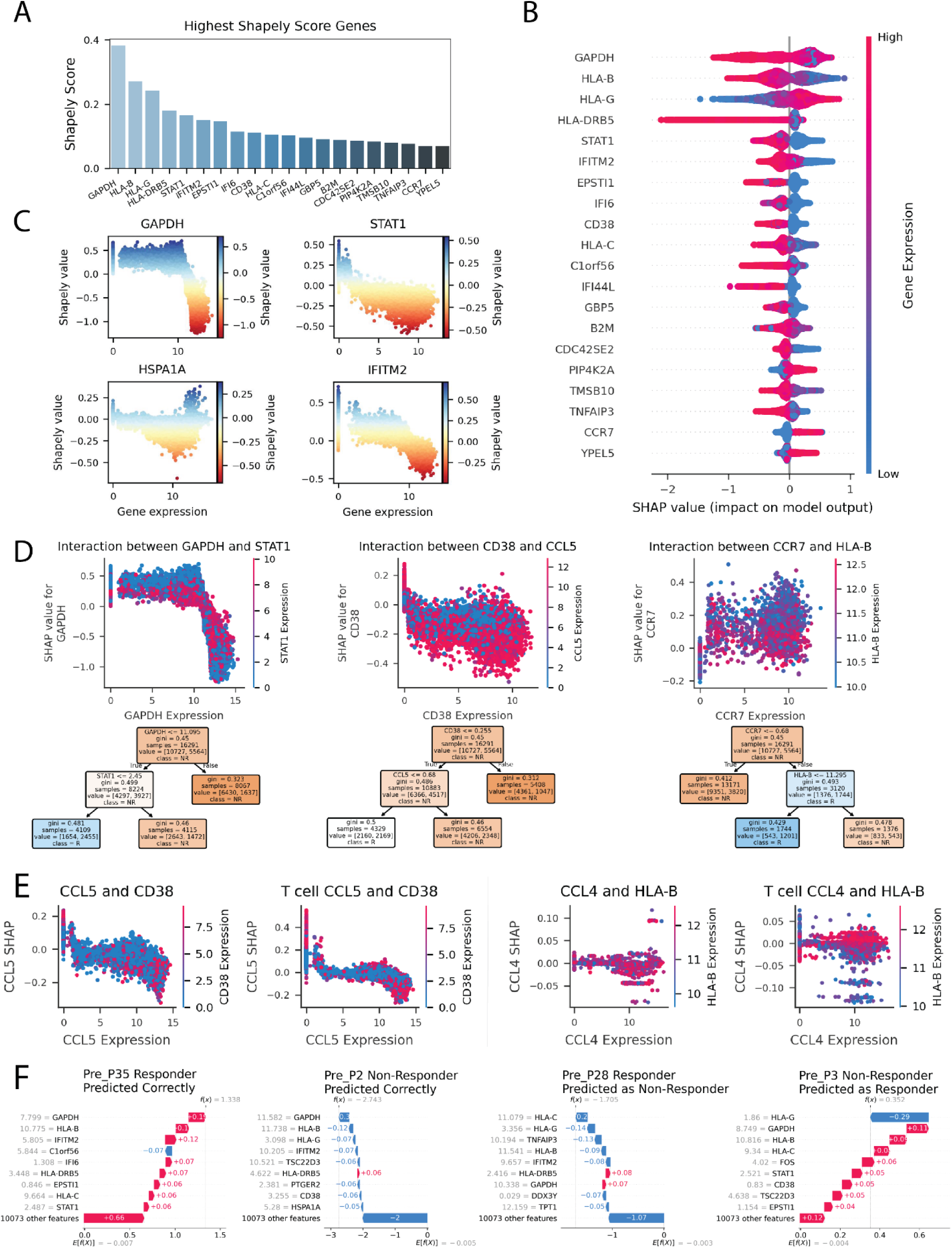
SHAP Analysis of Gene-Gene Interactions in Immune Response Prediction. **A** 20 Top genes with highest absolute Shapley score identified in the model. **B** SHAP value summary plot depicting the impact of each of the most important genes on model output. Gene expression (shown by coloring) as a function of SHAP value (x-axis) showing the relation between expression pattern and model’s prediction. **C** Scatter plots showing the relationship between gene expression and SHAP values for key genes (*GAPDH*, *STAT1*, *IFITM2*, and *HSPA1A*). **D** Interaction plots illustrating the SHAP value dependencies between specific gene pairs: *GAPDH* & *STAT1* (left), *CD38* & *CCL5* (middle), *HLA-B* & *CCR7* (right). Expression of the first gene shown as x-axis position, and the expression of the interacting gene shown as coloring; SHAP value shown as y-axis position. Decision trees beneath the plots show the simplified relation of the gene-pair conditional expression with the SHAP values, representing response to ICI. **E** Interaction plots comparing SHAP value dependencies between gene pairs trained on all cells vs T cells: *CCL5* & *CD38* (left), *CCL4* & *HLA-B* (right). Expression of the first gene shown as x-axis position, and the expression of the interacting gene shown as coloring; SHAP value shown as y-axis position. **F** Waterfall plots showing SHAP value contributions to individual model predictions (sample prediction). Four examples of patients predicted as responders or non-responders, with differences in gene expression patterns contributing to the outcome predictions. Pre_P28 – Responder predicted as Non-Responder, Pre_P3 – Non-Responder predicted as Responder, Pre_P2 – Non-Responder predicted correctly, Pre_P35 – Responder predicted correctly.

After training an XGBoost model over all of the cells in the cohort, SHAP values were calculated from that model, quantifying the contribution of each gene for each cell in the cohort. For example, as illustrated in Figure 2C, *GAPDH*, and *STAT1*’s expressions are associated with non-response. The directionality of the SHAP value is such that high SHAP values are associated with response, and low SHAP values with non-response. However, the relationships of the two genes differ: *GAPDH* exhibits a continuous relationship with a non-zero threshold separating responders from non-responders, whereas *STAT1* shows an on-off behavior, with zero expression linked to response and non-zero expression linked to non-response. Conversely, other genes such as *HLA-G* (Supplementary Figure 3) were associated with response. Additionally, SHAP reveals other, non-monotonic, gene behaviors. For example, *HSPA1A* (Figure 2C) shows a complex pattern where both very low and very high expression are associated with response, while the middle range is more related to non-response. Another interesting example is of *IFITM2* where zero expression is related to favorable response, the middle range has less effect on the prediction and high *IFITM2* pushes the model towards non-response.

Going beyond the effects of single genes, we next investigated SHAP values for gene interactions. Focusing on the top most important genes that were found in the FS section, we examined their most interactive partners in predicting immunotherapy response (Figure 2D). Notable pairs, showing a prominent conditional behavior in relation to SHAP, include *GAPDH-STAT1*, *HLA-B-CCR7*, and *CD38-CCR5* (additional pairs are shown in Supplementary Figure 3). To further simplify and enhance interpretability, we created decision trees for each gene pair. These trees align well with interaction plots, quantifying expression thresholds that best differentiate responses. As shown in Figure 2D, the interaction between *GAPDH* and *STAT1* highlights that cells with low *GAPDH* (<11.1) and high *STAT1* (>2.45) are associated with non-response, whereas low *GAPDH* with low *STAT1* is linked to response. High *GAPDH* is associated with response regardless of *STAT1* expression. The *CCR7-HLA-B* pair show a different relation to response, where the separation by *CCR7* is close to zero (<0.68), demonstrating an on-off conditionality, where low *CCR7* is associated with non-response. In higher values of *CCR7*, *HLA-B* best separates between outcomes with expression threshold of 11.3, where lower *HLA-B* expression relates to response, and higher to non-response. Such interaction analyses constitute a great way of modeling conditional effects of genes - relationships that are challenging to find without advanced models like SHAP.

SHAP dependencies for the model that is trained on all cells, captures in part the cell-types and sub cell-types expression patterns. Given that cells in different clusters exhibit distinct expression profiles, a cluster associated with response or non-response is likely to reflect its unique expression pattern in the dependence plot. For example, cluster G1 (B cells, n = 1445), which is significantly more abundant in responders, shows a nearly 2.5-fold higher proportion of low *GAPDH*, low *STAT1* cells compared to the entire cohort. In contrast, G6 and G11 (Exhaustion clusters, n = 2222 and n = 1129) are significantly more abundant in non-responders. Consistently, only a small fraction of these cells exhibits low *GAPDH* and *STAT1* expression, with G6 showing a 3.3-fold lower abundance, and G11 a 20-fold lower abundance compared to its proportion in the cohort. To explore intra-cell-type dependencies, we reduce the influence of this effect by also focusing solely on T cells (Figure 2E).

The T cell-specific analysis revealed that in certain gene pairs, interactions became more distinct. In particular, the separation of expression patterns was clearer in T cells for both *CCL5-CD38* and *CCL4-HLA-B* (Figure 2E), highlighting the importance of these interactions within this specific cell type. In the *CCL4-HLA-B*, the dependence is almost undetectable using the whole cohort, but with the T cells this is one of the strongest dependencies. In other gene-pairs, such as *GAPDH-STAT1*, the dependence is clearer in the whole cohort rather than in T cells, probably due to the effect of cell-type bound expressions described above.

Interestingly, all the chemokines in this analysis—*CCL4*, *CCL5*, and *CCR7*— have an on-off behavior. Non-zero expression, albeit low, of these chemokines is sufficient to dramatically alter the behavior of other genes in relation to the immune response (Figure 2D,E). *CCL4* and *CCL5* can promote anti-melanoma immune response (37), and are known to have a role in recruitment of immune cells (38,39). They have also been found to promote tumor development and progression by recruiting regulatory T cells to the tumor site (40). It is not surprising then, that their presence or absence can significantly change the effect of other genes on the immune response. This is especially true for *HLA-B*, which is critical for activating *CD8*^+^ T cells; the activation function can only be exerted following recruitment of these T cells, facilitated by chemokines like *CCL4*.

To further elucidate the behavior of the model, we leveraged SHAP values’ ability to deconvolve the contributing factors to each prediction, allowing us to understand the model’s choices. We analyzed the model’s performance to understand why certain samples were correctly or incorrectly predicted. We focused on four samples: Pre_P2 and Pre_P35 that were correctly predicted, and Pre_P3 and Pre_P28 that were misclassified. Using SHAP values, we trained the XGBoost model on all cells except those from the target sample, mimicking the actual prediction process. We then summed the contributions of all cells in each left-out-sample to obtain aggregated scores for each gene (Methods). These scores, sorted by absolute value, reveal the most critical genes for the model’s predictions (Figure 2F). SHAP waterfall plots are commonly used for interpreting a single prediction, but here, since each sample contains many prediction targets (cells), the plots represent an aggregation of these values. Therefore, the scores for each sample depict the average contribution of each gene to the prediction of all the cells in that sample – showing a general trend rather than a single contribution value. For instance, Pre_P35 was classified correctly as a responder, primarily due to low *GAPDH* and HLA-B values, with smaller contributions from several other genes. Pre_P2 was correctly classified as a non-responder, mostly due to a high *GAPDH* expression, as well as a high *HLA-B* and *IFITM2* expression and low *HLA-G* expression. Pre_P28 was misclassified as a non-responder due to the high *HLA-C*, *TNFAIP3*, and *HLA*-B expression, coupled with low *HLA-G*. Conversely, Pre_P3 was misclassified as a responder due to the average low expression of genes including *GAPDH*, *HLA-B*, *HLA-C*, *FOS*, *STAT1*, *CD38*, *TSC22D3*, and *EPSTI1*. Although a low *HLA-G* score pushed the model toward non-response, the cumulative effect of the rest of the genes was stronger.

Overall, the ability to interpret the model’s performance and decision-making is crucial for model improvement, gaining sample-level insights, and ensuring the model’s applicability in a clinical setting, which is the ultimate goal. Once the genes that affect the model’s prediction either positively or negatively are identified, it is possible to examine the distribution of these genes through the cohort and in the different cell-types.

### Reinforcement Learning Enables Labeling Each Cell’s Predictivity

Our analysis revealed that certain subsets of T cells perform best in predicting patient response. However, it remains unclear whether every cell within this group is necessary to achieve that level of predictive accuracy, or if some cells may not contribute to the model’s prediction and even detract from its performance. Specifically, up until now we considered all the cells from a responding sample as responders and all the cells from a non-responding sample as non-responders. This approach represents a simplified model rather than a reflection of reality, as the TME is heterogeneous, and contains synergistic, contradictory, and inert factors. For example, while the proportion or exhausted T cells mostly associates with non-responding samples in baseline, they do exist also in responders (41). To address that, we developed a novel reinforcement leaning (RL) framework that is built upon the basic model. Our goal was to quantify each cell’s effect on model prediction and create a cell-level score by rewarding cells with directional rewards when they are predicted correctly. By doing this, we move from a coarse, cluster-level measure, to a fine-grained, cell-level score.

First, to get a continuous score, the XGBoost model was switched from a Classifier model to a Regressor model. Moreover, to create the most accurate model possible, we only used the top genes from the FS section. This not only improved running time but also ensured a better distribution of the labels across samples. Initially, each cell from a responder sample is labeled 1, and each cell from a non-responder is labeled -1. Then, the model iteratively updates the continuous labels for each cell, ultimately assigning a score to each (Methods). Cells that are consistently predicted in the same direction will receive repeated rewards and therefore will be pushed to the edges of the distribution. In contrast, cells that are often predicted incorrectly will be iteratively normalized without any reward, resulting in scores with a low absolute value. These scores reflect the strength and direction of each cell’s contribution to the response of the sample (Figure 3A,B). Large positive and negative scores indicate strong predictive power of positive and negative response, respectively, while close to zero scores indicate an overall weak association of these cells with patient response.

**Figure 3:**
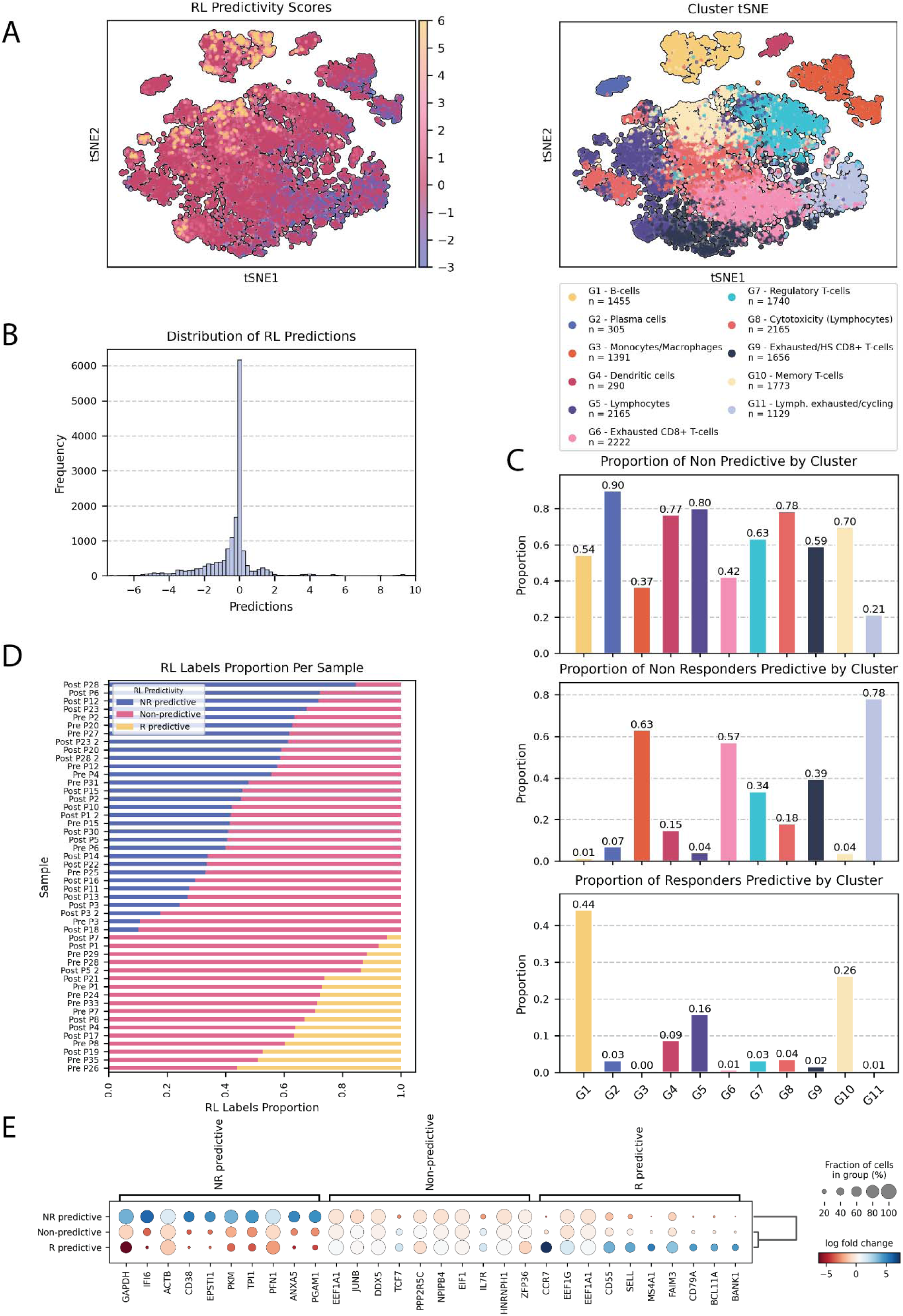
RL Prediction power. **A** tSNE plots showing immune cell clusters (top) and RL prediction scores (bottom). Clusters are colored by group, and the RL prediction scores range from highly predictive to non-predictive (lower limit is determined as -3 and upper limit as 6 for visualization purposes). **B** Histogram of RL prediction distribution. **C** Bar plots showing the proportion of non-responders predictive (top), non-predictive (middle), responders predictive (bottom) cells by cluster. **D** Stacked bar plots representing the proportion of RL labels (R predictive, non-predictive, NR predictive) for each patient sample. Top 31 samples are Non-Responders and bottom 17 samples are Responders. **E** Dot plot of the top most differentially expressed genes between the three RL bins, showing log fold change of key genes in non-responders predictive (NR predictive), non-predictive, and responders predictive (R predictive) groups.

Examining the score distribution across clusters and cell types, we categorized each cell into three bins based on its score: ‘non-response predictive’ (< -0.5), ‘non-predictive’ (between -0.5 and 0.5), and ‘response predictive’ (> 0.5). Figure 3A illustrates the distribution of these bins across the cohort. There is a clear association between the proportion and direction of predictive cells within each cluster and the relation of the cluster to response (Figure 3C), as determined by the original study (15). Specifically, clusters G3 (Monocytes/Macrophages), G6 (Exhausted CD8+ T-cells), and G11 (Lymphocytes exhausted/Cell cycle), which are prevalent in non-responders, also had the highest proportion of non-response predictive cells. Conversely, clusters G1 (B-cells) and G10 (Memory T-cells) that are associated with responders, were the top clusters in terms of response predictive cells. This analysis also highlights clusters G9 (Exhausted/HS *CD8*^+^ T-cells) and G7 (Regulatory T-cells), which have a relatively high prevalence of non-response predictive cells, and cluster G5 (Lymphocytes), which contains more response predictive cells, although with a lower proportion (see Supplementary Figure 5 for cell-type RL distribution). This observation serves as a sanity check, reinforcing the validity of our model.

Notably, non-predictive cells are well distributed across most clusters and are prevalent throughout the cohort. This might result from the arbitrary threshold used to bin the cells, however, examining the distribution of the scores across the cohort shows that most of the non-predictive cells are indeed close to zero and are therefore not affected by the specific threshold used (Figure 3B). In addition, it is important to note that these cells have passed rigorous quality control, so their non-predictive phenotype is not likely due to low-quality data but rather suggests a different level of participation in immunity. Finally, the spreading of non-predictive cells across different clusters and cell-types suggests that simply filtering cell-types or focusing on a single cluster is not sufficient, emphasizing the potential need for a finer method to filter cells and narrow the analysis to predictive cells.

Following, we examined the distribution of RL labels within each sample (Figure 3D). As expected, the proportion of predictive cells in well-classified samples was much higher, while misclassified samples had a higher proportion of non-predictive cells. Although likely reflecting to some extent the accuracy of the model, this also highlights the heterogeneity of the TME both within individual samples and across different samples.

To characterize these cells and understand what makes a cell predictive or non-predictive of patient response, we performed differential expression analysis among the three bins, as shown in Figure 3E. As expected, the genes used in the RL model appeared in the top differentiators list, but other significant genes also emerged, suggesting that the RL model manages to generalize and create biological groups. These include *PKM*, *PGAM1* and *ACTB* (42–44) in non-response predictive, *TCF7*, *IL7R* in non-predictive, and *SELL*, *MS4A1* (45) in response predictive cells. Although some biological genes were extracted as differentiators of non-predictive cells, this group does not seem to have strong markers differentiating it from the rest of the cells.

Quantifying individual cell participation in immune response provides novel, granular insights into the tumor microenvironment. This approach also offers an avenue for identifying and filtering out non-contributory cells, refining predictions related to immunotherapy response.

### Gene- and cell-based signatures predict patient response to ICI

To evaluate the generalizability of our findings, we first constructed a gene signature differentiating between responders and non-responders (Figure 4A). To this end, we extracted the genes with the highest feature importance scores from each fold of the LOO (Methods). Analyzing the intersection of genes from the 48 folds, we selected the top 11 genes that consistently intersected across the folds: *GZMH*, *LGALS1*, *GBP5*, *HLA-DRB5*, *CCR7*, *IFI6*, *GAPDH*, *HLA-B*, *EPSTI1*, *CD38*, and *STAT1*. Although somewhat arbitrary, the number of genes selected was based on the minimum number of top genes needed from each fold to form the intersection group: the top 11 genes required relatively few top genes from each fold, whereas those further down the list required a larger number of intersecting genes, making them less robust (Supplementary Figure 1). By examining the SHAP distribution for these genes’ expression, we determined that all except *CCR7* were negatively correlated with response. Following, the score we devised was an equally weighted combined score of the genes in the signature, with the weight of *CCR7* specifically negatively adjusted (Methods).

**Figure 4:**
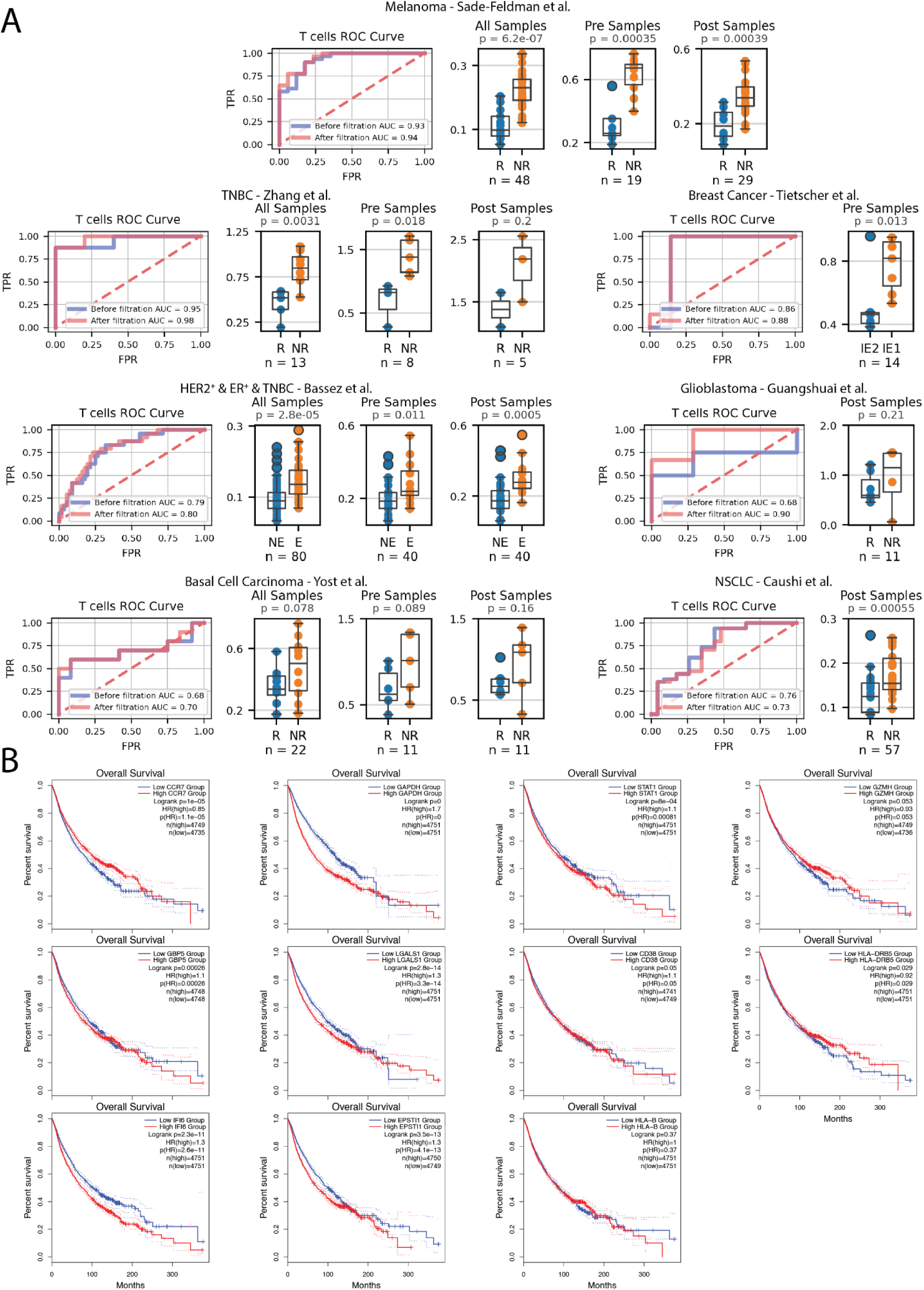
T Cell Response and Overall Survival Analysis in Various Cancer Types. **A** T Cell ROC curves showing the predictive power of the 11-gene signature score combined with RL-based filtration in distinguishing responders from non-responders across multiple cancer types. The blue curve represents the ROC curve before filtration, and the red curve represents the ROC curve after filtration. Box plots, separated by pre- and post-treatment samples (where available), depict the 11-gene signature scores before RL filtration. The datasets included are: Melanoma – Sade-Feldman et al. (15), TNBC (Triple-Negative Breast Cancer) - Zhang et al. (46), NSCLC (Non-Small Cell Lung Cancer) – Caushi et al. (47), BCC (Basal Cell Carcinoma) – Yost et al. (48), Glioblastoma – Mei et al. (49), HER2+ & ER+ & TNBC - Bassez et al. (50), Breast Cancer - Tietscher et al. (52). **B** Kaplan-Meier survival analysis of the 11 genes in the signature using bulk RNA-seq data across various cancer types. The survival curves illustrate the association between high and low expression of each gene. Plots include survival for genes – GAPDH, CD38, CCR7, HLA-DRB5, STAT1, GZMH, LGALS1, IFI6, EPSTI1, HLA-G, GBP5.

To mitigate biases arising from differences in cell-type abundance and considering that some datasets contain only T cells, we focused exclusively on T cells for all of our validation datasets. First, we evaluated the scores on the training melanoma cohort, which produced a high AUC score of 0.93 (n = 48, p-value = 6.16 × 10^-7^). Importantly, a significant difference was observed when examining baseline and post-therapy samples separately (n = 19 and 29, p-values of 3.5 * 10^-4^ and 3.9 * 10^-4^, respectively).

To establish the robustness of this score, we analyzed additional publicly available datasets having scRNA-seq data from patients treated with ICI (Figure 4A). In a triple-negative breast cancer (TNBC) dataset, our predictor achieved an AUC of 0.95 (n = 13, p value = 0.003) (46), and baseline samples were perfectly separated using the score receiving an AUC of 1 (n = 8, p-value = 0.018). In a non-small cell lung cancer (NSCLC) dataset containing only post-treatment samples, our score metric achieved an AUC of 0.76 (n = 57, p-value = 5.5 * 10^-4^) (47). Two additional datasets showed a positive trend in association with the score, though they were not as significant with this simple score: a glioblastoma dataset and a Basal-cell carcinoma (BCC) dataset both receiving an AUC score or 0.68 (n = 11, p-value = 0.2 and n = 22, p-value = 0.08, respectively) (48), (49). The BCC dataset showed a relatively better separation using only baseline samples (AUC = 0.77), though due to the smaller sample size, this separation remained insignificant (n = 11, p-value = 0.09).

Next, we applied our predictor to a dataset of HER2 positive, ER positive and TNBC patients having annotations for clonal expansion phenotype (50). The dataset consists of two cohorts, one of treatment naive samples, and the second of patients receiving neoadjuvant chemotherapy prior to anti-PD1. In both cohorts, the score was significantly associated with expansion, with the first cohort receiving an AUC of 0.81 and the second 0.77 (n=58, p-value = 8.1 * 10^-5^, and n=22, p-value = 0.03). Combined, these datasets achieved an AUC score of 0.79 (n = 80, p-value = 2.8 * 10^-5^). This trend remained significant when separating baseline and post-treatment samples. This finding suggests that samples with expanded clonal populations are more likely to be non-responsive. This also aligns with a study in melanoma patients, which showed that highly expanded clonotype families were predominantly distributed in cells with an exhausted phenotype, characterized by decreased diversity of T cell receptors (TCRs) (51). Lastly, we applied our signature to a dataset of breast cancer patients having a classification for exhausted and non-exhausted TME (52). This data predominantly contained samples from treatment-naive breast cancer patients. Our predictor achieved an AUC of 0.86 and significantly differentiated between these two TME states (n = 14, p value = 0.01).

Following, we sought to validate the RL based filtration of non-predictive cells by constructing a cell-filtration score (Methods). This score guided the removal of cells from each dataset, after which we reapplied the 11-gene signature to the remaining cells and tested for prediction accuracy (Figure 4A). While the genes identified in the differential expression analysis in the previous section effectively differentiate response-predictive from non-response-predictive cells, as aforementioned, non-predictive cells had less obvious distinguishing markers (Figure 3E). Since we aim to apply the score to remove non-predictive cells in new datasets, a more sophisticated way of characterizing these cells is needed. To address this, we used a logistic regression model to predict who are the non-predictive cells. Although this classifier cannot be directly applied to other datasets due to differences in scale and potential batch effects, we can derive a more generalizable score from it that can be applied to new data. The filtration score was built according to the logistic regression classifier of non-predictive cells, using the coefficients of the top 100 most important genes (Supplementary Table 5). A weighted mean over all these genes determined the RL-based filtration-score (Methods).

To balance the removal of non-predictive cells while minimizing the risk of filtering out predictive ones, we chose to remove 40% of the cells with the highest scores across all datasets. The results were robust to the threshold choice between 15-60%: as expected, less filtration led to smaller improvements, while more filtration resulted in greater improvements, up to a certain point (Supplementary Table 6). In the melanoma dataset that was used for devising the gene and cell scores (15), the addition of the cell-filtration improved the AUC score to over 0.94 (n = 48, p-value = 2.55 × 10^-7^). Applying this filtering to the other datasets used for validation, achieved an improvement in the prediction accuracy in five out of the six. Specifically, the glioblastoma dataset showed a significant improvement in AUC, from 0.68 to over 0.9 (n = 11, p-value = 0.03). Filtration in the BCC dataset produced an improved AUC of 0.7 (n = 22, p-value = 0.06). The HER2 positive ER positive and TNBC dataset consisting of the two cohorts, saw an increase in AUC to 0.8 in relation to expansion (n = 80, p-value = 1.3 * 10^-5^). The AUC of the breast cancer cohort labeled with exhaustion increased to 0.88 (n = 14, p-value = 0.009). In the TNBC (46) the AUC was improved from 0.95 to 0.98 (n = 13, p value = 0.0016). In the NSCLC dataset the AUC score was reduced to 0.73.

### Gene signature predicts patients’ survival

Finally, we examined the specific genes that make up the 11-gene signature, using bulk RNA-seq datasets to assess their association with patient overall survival (Figure 4B). We expected that higher immune responsiveness would be associated with better survival, hence the signature should differentiate survival rates in this direction. Of note, validation in bulk RNA sequencing is challenging due to the mixture of different cell types. However, since survival analyses look at outcomes across a cohort, they are less dependent on the precision of cell-type-specific resolution. What matters is whether the overall expression of these immune genes, in the context of the mixed population, correlates with better or worse outcomes for the patients. Moreover, strong immune-related signals can still emerge because immune gene expression often reflects the overall immune activity in the tumor microenvironment.

To examine this, we used the GEPIA2 online tool (53), and assessed the relation between each of the 11 genes and survival. Using the median expression as a cutoff and using all cancer types available in GEPIA (Methods), we found that 7 out of the 11 genes, *CCR7*, *STAT1*, *GBP5*, *LGALS1*, *EPSTI1*, *IFI6* and *GAPDH*, were significantly differentiating the overall survival time in the expected direction (after Bonferroni correction). The remaining 4 genes, *HLA-DRB5*, *HLA-B*, *GZMH* and *CD38*, showed no significant association with survival in either direction (Figure 4B). The fact that this signal comes from all cancer types and shows that patients with high expression of these genes in their tumor exhibit an improved survival rate, strengthens our hypothesis that this could indeed be the immune related activation.

## Discussion

In this study, we developed PRECISE, a machine learning framework designed to predict response to ICI from single-cell RNA-seq data. Our aim was to provide a roadmap for how single-cell data can be used in prediction tasks, such as treatment response, while maintaining model interpretability. Machine learning models stand out as the only technology able to capture the complexity level seen in the TME and reflected in single-cell data. The main advantage of these supervised learning models is that they are trained directly on the target and adapt to the characteristics of the data, instead of merely searching for patterns linked to the target.

The ability to work simultaneously on two levels of complexity—sample-level and cell-level—is crucial for any model designed for single-cell interpretation. Our model enables exactly that: initially, by assigning each cell a label according to its sample; next, by combining cell-level predictions into a sample response score; later, by aggregating the single-cell effects on the model using SHAP and grouping them into sample contributions; and lastly, through the reinforcement learning model, diffusing the sample responses back down to assign a predictivity score to each cell.

Our model is highly flexible, both in terms of implementation and objective. Although we used XGBoost, many other algorithms with sufficient capacity could handle the data complexity equally effectively. Additionally, we opted to include all genes that passed preprocessing steps, capturing the strongest signals across the data. However, this approach may overlook significant subsets of the data. Future analyses could focus on specific gene groups (e.g., metabolic genes) to explore different dimensions and revealing new insights.

Differences between cancer types, as well as the differences in prior treatments and medical backgrounds, make transferability of results from one dataset to another very challenging. Despite these challenges, the 11-gene signature showed a positive trend in all datasets, indicating the robustness and generalizability of the model’s results. The RL filtration score also showed a positive trend in most datasets, improving prediction accuracy and effectively removing noisy, non-predictive cells.

Focusing on the immune system is an excellent way to establish transferability between cancer types, since while the cancers can greatly differ, the immune system retains common features. This likely explains why our model’s results could be generalized across different cancer types. However, the model was only trained on one dataset. Training on multiple datasets together, or transferring the model directly to new datasets was infeasible due to the large differences in expression between cohorts and especially across sequencing technologies. This limits the model’s scalability and its ability to generalize. Future work should aim to address these limitations, and the best current approach may be to use foundation models that integrate single-cell data into a unified embedding (54,55) (interpretability should be kept in mind).

Incorporating multiple datasets from different cancer types will significantly enhance the robustness of such models. Scaling up the size of the data, and number of samples used, will open additional possibilities, such as adding higher-level models that aggregate scores from different cell types to produce an overall sample score.

Another limitation is the use of both baseline and post-treatment samples. While the ultimate objective of such models is the prediction of patient response before treatment, we included in our training data both pre- and post-treatment samples to maintain as much data as possible for the training step. Nonetheless, we found that the prediction accuracy was similar in the validation datasets when estimated on pre-treatment samples alone, and in some cases even produced better separation. Moreover, it should be noted that prediction on post-treatment samples has a value in cases of advanced line ICI re-challenge, for patients not responding to first line treatments.

## Conclusions

This study demonstrates the potential of machine learning models trained on single-cell data to predict immunotherapy responses and provide deep biological insights. Using PRECISE (Predicting therapy Response through Extraction of Cells and genes from Immune Single-cell Expression data), we offer a roadmap through which single-cell data can be used not only for sample predictions, but also for understanding complex biological systems such as the TME. Machine learning models in this context can go beyond mere prediction – we showed their utility in capturing gene-gene interactions, identifying non-linear expression patterns, and quantifying cell-type and single-cell participation in immune response. With sufficient data and further development, these models could become exceedingly powerful tools. Expanding training to larger, multi-dataset cohorts will be crucial for improving generalization.

## Methods

### Data Collection and Preprocessing

We utilized a publicly available scRNA-seq dataset containing immune cells from melanoma patients, generated using Smart-seq technology (15,56). The dataset included 16,291 immune cells from 48 samples. Following initial quality control done by the authors, preprocessing primarily involved filtration of genes based on expression levels. Specifically, we excluded non-coding genes, mitochondrial genes, and ribosomal protein genes (genes starting with ‘MT-‘, ‘RPS’, ‘RPL’, ‘MRP’, and ‘MTRNR’). Only genes expressed in at least 3% of cells were retained, resulting in 10,082 genes. Cells were labeled according to their sample’s response to ICI therapy, with cells from responding samples labeled as responders and those from non-responding samples labeled as non-responders.

Additional datasets were collected and preprocessed for validation, including droplet-based datasets containing read counts (46–50). In the droplet-based datasets, expression levels were first normalized to a standard sum of 10,000 reads per cell and then log-transformed. It is important to note that the per-cell read coverage in these datasets was often higher than 10,000. Only samples with defined clinical annotations (response or clonal expansion) were included in the analysis. We considered only patients treated with either immune checkpoint inhibitors (ICI) or a combination of ICI and chemotherapy. Additionally, a dataset of a non-ICI cohort containing mostly treatment-naive patients with annotations for exhausted and non-exhausted tumor microenvironment (TME) was included (52).

### Cell-Type Assignment and Unsupervised Clustering

In our previously published melanoma dataset (15), we used the original classification of T cells that was based both on gene markers and manual curation. Eleven unsupervised clusters and supervised cell-type assignments of the cells were obtained from the original article. In most other datasets, T cells were either extracted using flow cytometry, or were identified *in silico* using computational approaches in the paper (47–50,52) generating the data. The TNBC dataset (46) was the only dataset without cell-type annotations. Identification of T cells in this dataset was done by unsupervised clustering, followed by differential expression on these clusters and determination of T cells according to known markers.

### Base Machine Learning Model

Training was conducted in a leave-one-out (LOO) cross-validation manner, where the model was trained on all samples except one, and predictions were made for the left-out sample. The model was trained and used for prediction at the single-cell level, learning to predict the label associated with the sample’s response. After predicting the labels of all cells in the cohort, we calculated the proportion of cells predicted as responders in each sample, forming the sample score. Accuracy of the prediction was evaluated using the Receiver Operating Characteristic (ROC) Area Under the Curve (AUC) score between the sample scores and the sample response labels, making the accuracy assessment robust to the threshold choice. From each model fold, feature importance scores were extracted for each gene using the model’s built-in ‘feature_importance_’ attribute, which quantifies each gene’s contribution to the model’s prediction.

### XGBoost Classifier Supervised Predictive Model

An XGBoost model, known for its balance between accuracy and interpretability, was trained to predict the response label of each cell (16). The XGBoost model was built using the XGBoost Python package. The classifier was trained using the default parameters for XGBoost, except for a learning rate (LR) of 0.2, a max_depth of 7, and an alpha of 1. The model’s performance was not sensitive to these parameters. No hyperparameter tuning was required. The objective function was set to ‘binary:logistic’. An important parameter included was the ‘scale_pos_weight’, set to the ratio of the number of negative cells to positive cells, addressing potential bias towards responders or non-responders in the data. Although accuracy was high without it, including this parameter ensures better generalization for more biased data.

### Per Cluster/Cell-Type Prediction

Cellular groups we’re extracted from the original paper (15), and included 11 clusters determined by unsupervised clustering, and 13 cell types or states determined using known gene markers. To examine cluster- and cell type-specific predictions, we ran the classification model with LOO on each cellular group, and obtained an AUC score for each cluster independently. Samples were excluded from the leave-one-out process if they didn’t contain any cells from the specific cellular group, and the accuracy scores were adjusted accordingly.

### Boruta Feature Selection

We used the boruta_py from the Python Boruta package. Boruta is a wrapper algorithm that iteratively removes features that are statistically less important than random probes (shadow features). We used Boruta to extract only the confirmed genes from its output in a leave-one-out fashion. We set alpha to 0.02, which produced a small, robust number of features. The chosen features were saved for interpretation and used for LOO prediction, ensuring no train-test leakage.

### Lasso Regression Based Feature Selection

Lasso regression, a technique that performs both variable selection and regularization, was used for feature selection. Lasso zeroes the coefficients of non-contributing features. Using different values of the regularization parameter alpha, we performed LOO predictions with Lasso-based feature selection. Although results were reasonable, they were consistently lower than those obtained with Boruta and required handpicking of the alpha value. Feature selection was done after scaling the genes, using Lasso from Python’s sklearn package (with max_iter set to 500).

### Simple Feature Selections

We also examined the performance of simple feature selection methods, such as differentiating genes and the most highly variable genes. Differential expression analysis between responding and non-responding samples was conducted within the LOO using either Wilcoxon ranksum test, t-test (scanpy), or Fisher’s exact tests with significance values of 0.05, 0.01 and 0.001. Highly variable genes were extracted using scanpy’s function on the top 1000, 2000, 4000 and 8000 top highly variable genes. The Wilcoxon ranksum test produced the best results among these methods but still yielded consistently lower performance compared to Boruta and resulted in a much higher number of genes.

### SHAP Analysis

SHAP (SHapley Additive exPlanations) values were used to interpret the model’s predictions. SHAP values explain the impact of each feature (gene) on the model’s output, providing insights into gene importance and interactions. We applied SHAP using the SHAP Python package on the genes: [*‘GZMH‘, ‘LGALS1‘, ‘GBP5‘, ‘HLA-DRB5‘, ‘CCR7‘, ‘IFI6‘, ‘GAPDH‘, ‘HLA-B‘, ‘EPSTI1’, ‘CD38’, ‘STAT1’*]. SHAP automatically identified gene interactions and interpret the mean contributions of genes to a sample’s prediction. We averaged the SHAP scores for each gene across the sample to generate sample-level Waterfall plots (Figure 2, Supplementary Figure 4).

### Decision Trees for SHAP Interaction Analysis

To model the pairwise conditional behaviors of genes as extracted by SHAP analysis, we used decision trees. Each time, we insert the expression pattern of two genes extracted by SHAP to the decision tree model. We used the DecisionTreeClassifier from python’s sklearn with max_depth set to 2. This limit on the depth helped exhibit a simple relation between the two genes.

### Reinforcement Learning Framework

To quantify the directional predictivity of each cell, we constructed a reinforcement learning framework based on XGBoost, but this time we used a regression model. The initial labels were set to 1 for cells from responders and -1 for cells from non-responders. Iteratively, we updated the labels of the cells if they were predicted correctly, weighted by the prediction value. Each iteration normalized the absolute value of the labels to keep their scale consistent. The pseudocode below illustrates this process:

<u>Initialization:</u>

1. Initialize responder cells with label 1 and non-responder cells with label -1
2. Set the list of chosen features to those extracted with feature selection
3. Set the learning rate (eta = 0.01) and the number of iterations (200)

<u>Iteration Process:</u>

For each iteration:

1. For each sample:

a. Train the model on all samples except the current sample.
b. Predict the labels for the current sample.
c. for each cell in sample:

- If the sample is a non-responder and the prediction value is negative, or if the sample is responder and the prediction value is positive:
  - Update the label: label = label +eta*prediction
  - Else:
  - Do not update the label.
2. Normalize the absolute updated value of the labels to maintain their scale.

Return:

Return the updated labels.

The labels were then classified into ‘Responder Predictive’ for *label >* 0.5, ‘Non-Responder Predictive’ for *label* < -0.5, and ‘Non-Predictive’ for -0.5 < *label* < 0.5. These groups were then characterized with differential expression analysis.

### Application of 11-gene Signature on External Datasets

We created a pseudo-bulk expression matrix by averaging the expression of each gene across all T cells in each sample. To normalize the differences in the scale of expression among different genes, we divided the expression of each gene in a sample by the total expression of that gene across all samples. Formally,

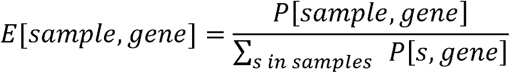

Where P is the pseudo-bulk expression matrix with the samples as rows and genes as columns, and E is the new, scaled matrix. The normalization term in the denominator sums the expression of a given gene over all samples in the dataset.

After calculating the scaled pseudo-bulk, the score for each sample was calculated as follows:

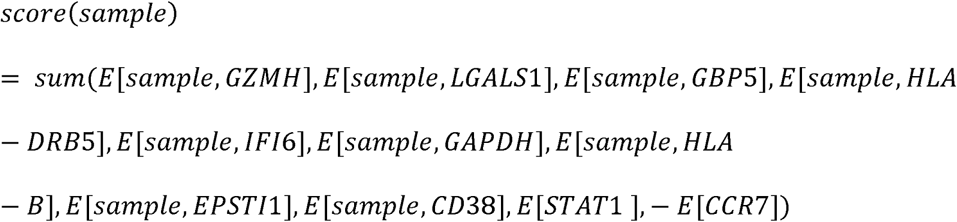

This score was then used to differentiate between the different responses in the samples.

### Cell Filtration Score Calculation and Application

The filtration score was calculated based on a logistic regression model. The model was trained to differentiate non-predictive cells from the rest of the cells (response predictive and non-response predictive) using the python LogisticRegression algorithm from sklearn (57), with the max_iter parameter increased to 500 to assure convergence. The 100 genes with the largest absolute coefficient values were extracted from the model and used to create a weighted score. Given the vector of the weights extracted by the logistic regressor and the expression matrix E normalized for each gene’s expression:

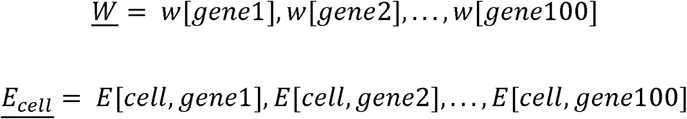

The score for a specific cell is calculated by the dot product of these two vectors

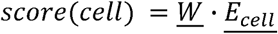

The cells with the largest scores are most likely to be non-predictive. The cells with the top 40% scores were filtered out in all datasets.

### Bulk RNA-seq Survival Analysis

GEPIA2, an online tool for analyzing RNA sequencing expression data, was used to examine the relation of the expression of the 11 genes to overall survival (53). Using the median expression as a cutoff, this produced Kaplan-Meier survival curves for each gene, over all available cancer types in GEPIA2. These include: ACC, BLCA, BRCA, CESC, CHOL, COAD, DLBC, ESCA, GBM, HNSC, KICH, KIRC, KIRP, LAML, LGG, LIHC, LUAD, LUSC, MESO, OV, PAAD, PCPG, PRAD, READ, SARC, SKCM, STAD, TGCT, THCA, THYM, UCEC, UCS and UVM.

## Supporting information

Description of Supplemental Tables

## Abbreviations

scRNA-seq: Single-cell RNA sequencing
TME: Tumor microenvironment
ICI: Immune checkpoint inhibitors
LOO: Leave-one-out
ROC: Receiver Operating Characteristic
AUC: Area Under the Curve
SHAP: SHapley Additive exPlanations
RL: Reinforcement learning
FS: Feature selection
HNSCC: Head and neck squamous cell carcinoma
BCC: Basal-cell carcinoma
NSCLC: Non-small cell lung cancer
TNBC: Triple-negative breast cancer
ADCC: Antibody-dependent cellular cytotoxicity
ML: Machine learning
NR: Non-responder/Non-responsive
R: Responder/Responsive
LR: Learning rate
HVG: Highly variable genes
TCRs: T cell receptors

## Declarations

### Ethics approval and consent to participate

Not applicable.

### Consent for publication

Not applicable.

### Availability of Data and Materials

All data used in this study is published and publicly available. The scRNA-seq data for the melanoma training cohort used in this paper is available through accession number GEO: GSE120575 (15). TNBC scRNA-seq data was downloaded from <u>GSE169246</u> (46). NSCLC scRNA-seq data was downloaded from <u>GSE176021</u> (47). Glioblastoma scRNA-seq was downloaded from Figshare (https://doi.org/10.6084/m9.figshare.22434341) (49). BCC was attained from <u>GSE123813</u> (48). HER2 positive, ER positive and TNBC scRNA-seq data was accessed from the Lambrecht’s lab website (https://lambrechtslab.sites.vib.be/en/single-cell) (50). Breast cancer scRNA-seq data classified by exhaustion was obtained from ArrayExpress database at EMBL-EBI under <u>E-MTAB-10607</u> (52). The methods described in this manuscript have been implemented in a tool called PRECISE, which is freely available at the GitHub repository. The repository also includes a tutorial for its use: https://github.com/yizhak-lab-ccg/PRECISE-.

### Author’s Contribution

K.Y. supervised the project and guided it throughout. K.Y. and A.P. jointly wrote the main manuscript text. A.P. prepared the figures and performed the analysis. Both authors reviewed and approved the final manuscript.

## Acknowledgements

We would like to thank Prof. Gad Getz, Dr. Moshe Sade-Feldman, Dr. Liron Zisman, Elad Zisman, Ofir Shorer, and Sapir Levin for fruitful discussions and helpful comments on the manuscript. Figure 1A was created with BioRender.com using a paid license.

