## Supplementary material for "Uncovering Predictive Gene and Cellular Signatures for Checkpoint Immunotherapy Response through Machine Learning Analysis of Immune Single-Cell RNA-seq Data": Description of Supplemental Tables

SuppTable1 - ModelScores
Consists of two sheets: BasicModelScores and BorutaModelScores. Each sheet provides the scores produced by the respective model for each sample, per cellular group, along with a Response column (1 for responder, 0 for non-responder).

SuppTable2 - MeanFeatureImportance
Maps the mean feature importance score for each feature across all folds in the LOO process of the base model (all cells with all genes included).

SuppTable3 - FS_scores
Consists of two sheets: SimpleFSAUCs and LassoAUCs. Each sheet contains a per-cellular group AUC score for the hyperparameters in each feature selection method.

SuppTable4 - Boruta_intersectionCount
Lists the number of times each gene was selected by Boruta across all LOO process iterations, per cellular group.

SuppTable5 - RL_non_predictive_LogReg_coeffs
Maps each of the top 100 genes to its coefficient as determined by the logistic regression model for predicting non-predictive cells.

SuppTable6 - AUC_scores_validation_RL_filtration
Contains the AUC scores of the 11-gene signature after RL-based cell filtration. For each dataset, the table lists AUC scores following filtration at various cell filtration percentages: 15, 20, 25, 30, 35, 40, 45, 50, 55, and 60%.
