## Supplemental Figures for "Uncovering Predictive Gene and Cellular Signatures for Checkpoint Immunotherapy Response through Machine Learning Analysis of Immune Single-Cell RNA-seq Data"

#### Supplementary Figure 1

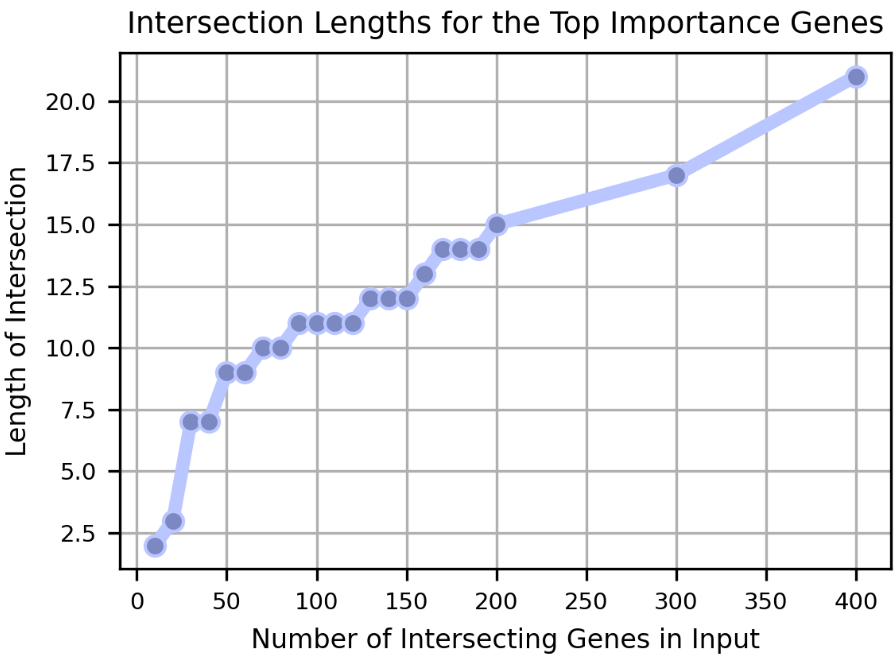


**Supplementary Figure 1:**
The plot shows the relationship between the number of genes in the output intersection group (y-axis) and the number of genes used as input (x-axis) in a feature importance-based vfeature selection (FS) process.

### Supplementary Figure 2

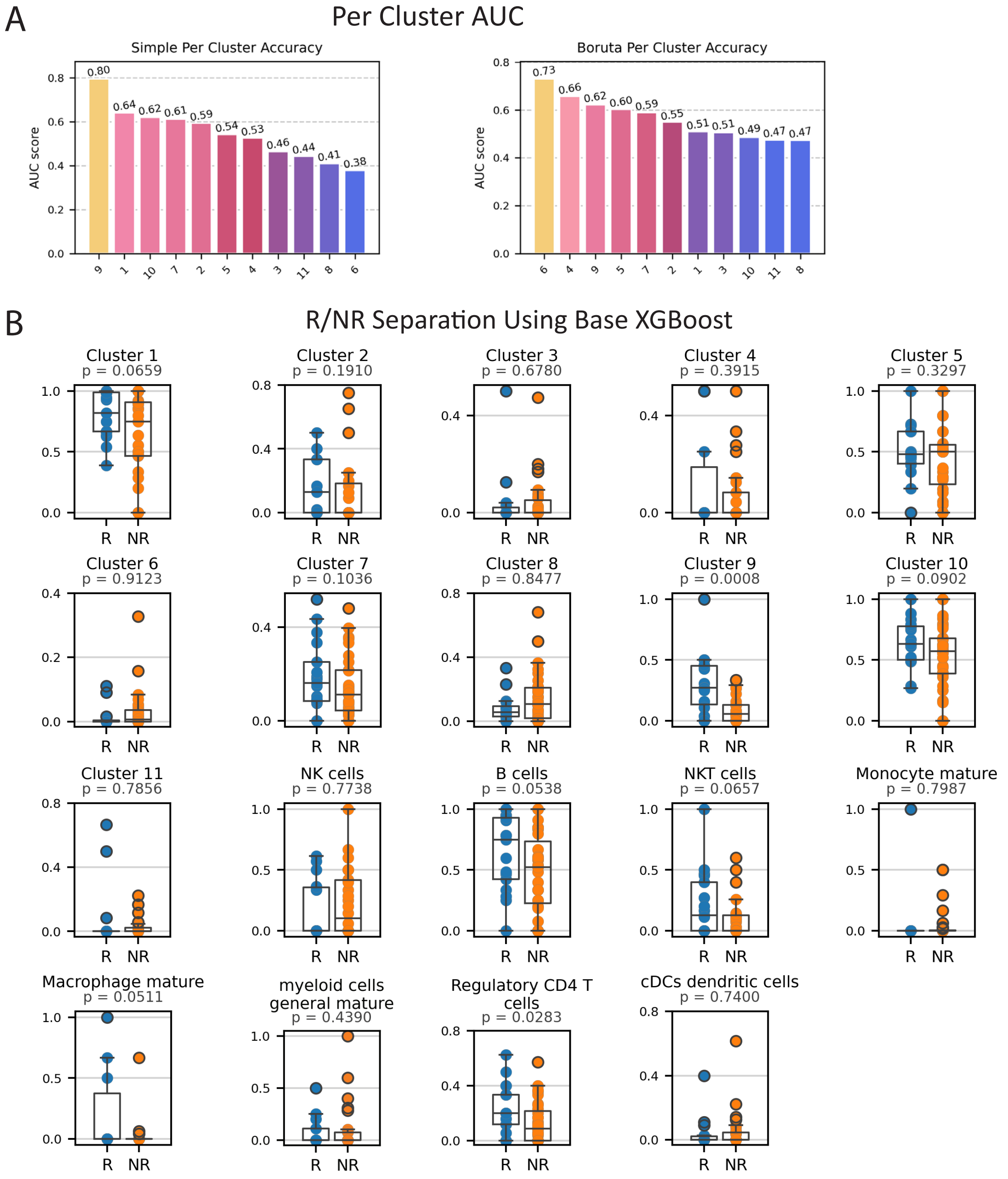


**Supplementary Figure 2:**

**A** Per Cluster AUC (top) - AUC scores indicating the prediction accuracy of the base model (left) and Boruta model (right) across the different clusters. **B** R/NR Separation Using Base XGBoost (bottom) - Box plots comparing the scores produced by base XGBoost between responders (R) and non-responders (NR) across cell subtypes and clusters, with p-values indicated by the Mann–Whitney U test.

### Supplementry Figure 3

## A

SHAP Values - Shap to Expression in Chosen Genes

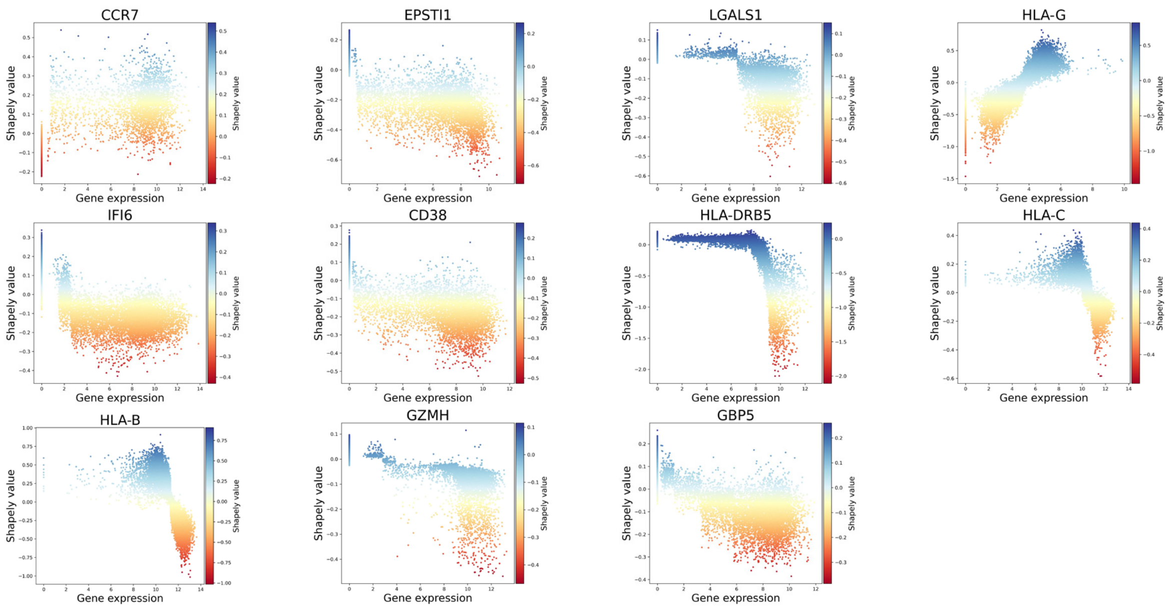


## B

SHAP Values - Gene-Gene Interactions

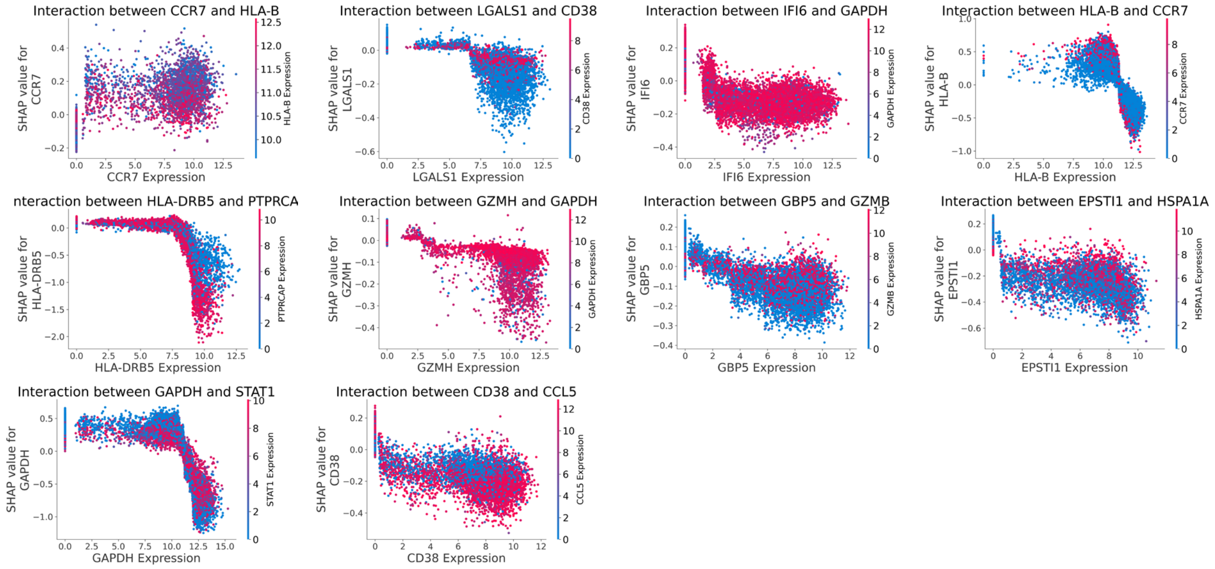


**Supplementary Figure 3:**
**A** Shap to Expression in Chosen Genes (top) - Scatter plots showing the relationship between gene expression and SHAP values for key genes. **B** Shap Gene-Gene Interactions (bottom) - Interaction plots illustrating the SHAP value dependencies between 11 genes from signature, and their most interactive pairs.

### Supplementry Figure 4

#### Waterfall Plots Per Sample - Interpretation of the Predictions

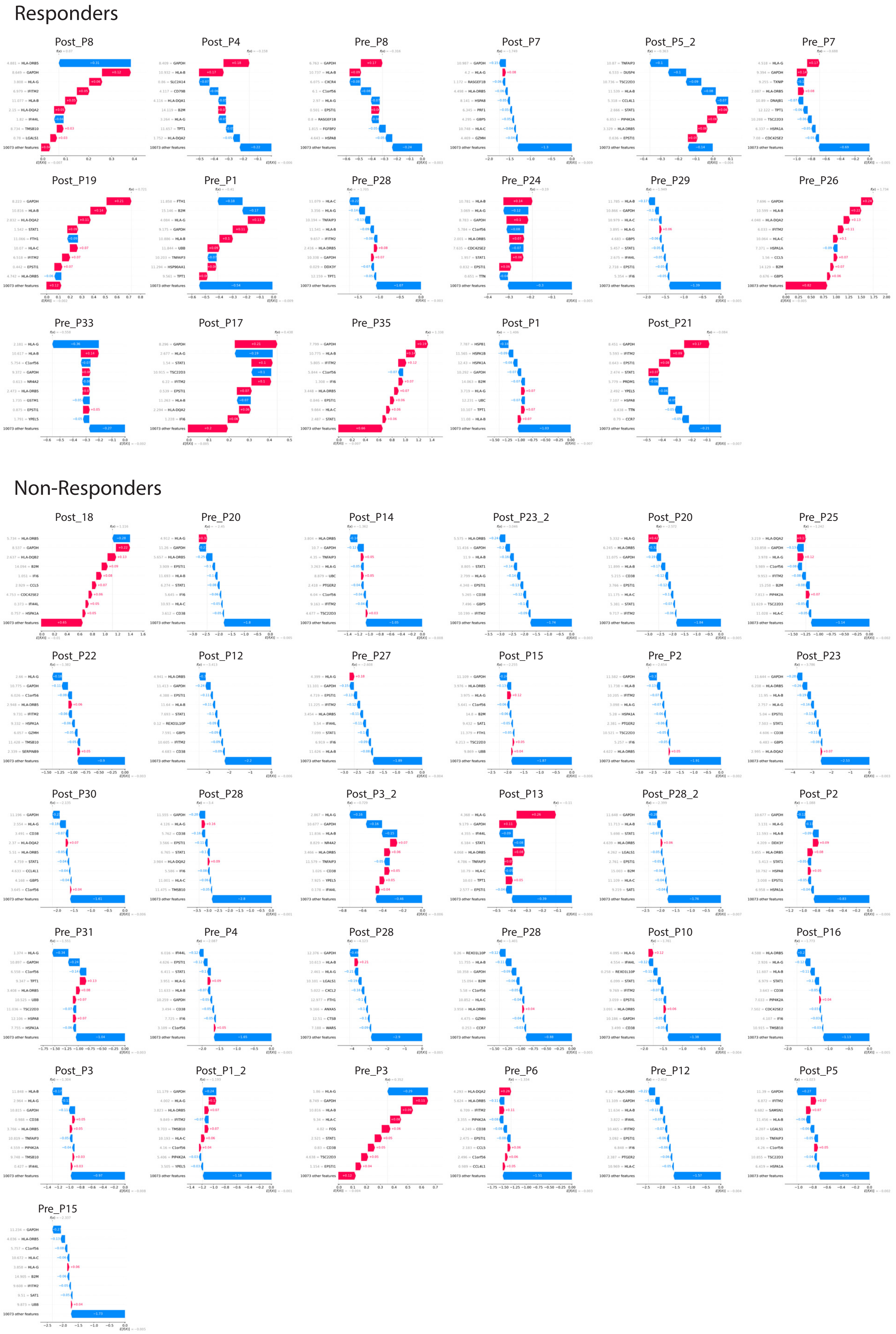


**Supplementary Figure 4:**
Waterfall Plots Per Sample - Waterfall plots showing SHAP value contributions to model predictions (sample prediction), separated to Responders (top) and non-Responders (bottom).

#### Supplementry Figure 5

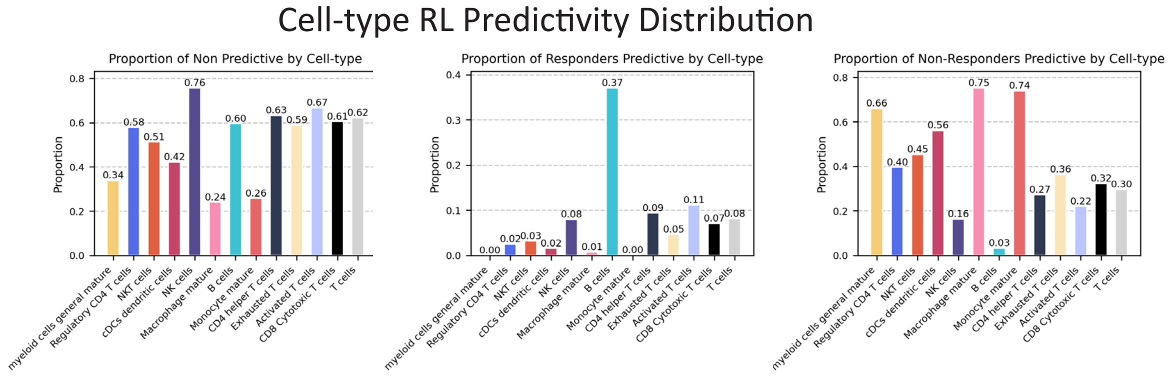


**Supplementary Figure 5:**
Cell-type RL Predictivity Distribution - Bar plots showing the proportion of non-responders predictive (left), responders predictive (middle), non-predictive (right), cells by cell-types.

#### Supplementry Figure 6

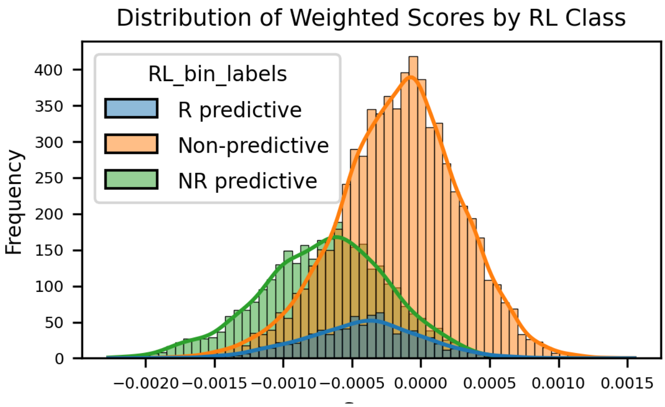


**Supplementary Figure 6:**
Distribution of weighted Scores by RL Class - Distribution of non-response weighted scores for RL prediction across the three RL bins – response predictive (blue), non-predictive (green) and non-response predictive (orange).
